# Proteasome inhibition enhances latency reversal and boosts NK cell-mediated elimination of HIV-1 infected cells through HLA-E downregulation

**DOI:** 10.1101/2024.10.18.619103

**Authors:** Thessa Laeremans, Sabine den Roover, Stefan Johan Nezic, Sabine D. Allard, Joeri L. Aerts

## Abstract

The shock and kill strategy primarily depends on using latency reversal agents (LRAs) to reactivate the dormant viral reservoir, rendering it visible for recognition and subsequent elimination by the host’s immune system. While this approach has shown high efficacy *in vitro* and *ex vivo*, its *in vivo* application has yet to show significant delays in time to viral rebound. This lack of *in vivo* efficacy is most likely due to the insufficient elimination of reactivated reservoir cells by the host’s immune effector cells, including natural killer (NK) cells. Given the pivotal role of NK cells in antiviral immune responses, we hypothesized that they are crucial players in pursuing a functional cure against HIV-1. However, the inhibitory interaction between NKG2A and HLA-E diminishes their effectiveness. Notably, proteasome inhibition has been effective in reducing HLA-E expression on various tumor cell types, thereby enhancing NK-cell mediated killing. However, its impact on HIV-1 latency remains unexplored. We found that the proteasome inhibition could reverse the latent state of J-Lat cells while substantially reducing HLA-E expression. Additionally, a reduced expression of NKGA on primary NK cells was observed which led to an increase in NK-cell cytotoxicity. These results suggest that disrupting the NKG2A/HLA-E interaction could potentially augment the effectiveness of the shock and kill strategy by improving NK cell-mediated clearance of reactivated cells.

**Importance:** Despite promising *in vitro* results, purging the viral reservoir using LRAs has yet to demonstrate clinical benefits. A significant challenge lies in the inadequate activation of immune effector cells, such as CD8^+^ T cells and NK cells. Therefore, developing therapeutic strategies to address these challenges could enhance the effectiveness of the shock and kill strategy. This study highlights the need for therapeutic interventions to overcome these hurdles. Our findings show that proteasome inhibition not only triggers latency reversal but also enhances NK-cell mediated elimination of latently infected cells *in vitro* by downregulating HLA-E. This suggests that targeting the proteasome could be a novel therapeutic approach in the shock and kill strategy, potentially improving clinical outcomes.

## Introduction

Although antiretroviral treatment (ART) effectively suppresses human immunodeficiency virus (HIV) replication, the existence of a viral reservoir prevents complete eradication of the virus. As a result, lifelong adherence to ART is required, as rapid viral rebound occurs almost invariably upon treatment interruption. To circumvent this dependence on ART, researchers are working towards a functional cure for HIV-1, where viral replication is controlled without the need for continuous ART and without complete viral eradication. Various functional cure strategies have been explored, focusing either on boosting the immune system, e.g. using therapeutic vaccination, or directly targeting the viral reservoir, as with the shock and kill strategy. The shock and kill strategy aims to purge the latent reservoir, allowing immune effector cells to eliminate the reactivated virus while maintaining ART. Although this strategy has shown auspicious results *in vitro* and *ex vivo*, clinical trials have not yet demonstrated significant delays in viral rebound. This discrepancy is likely due to the highly diverse nature of the viral reservoir and inadequate immune cell activation *in vivo* (1–3). While the latency-reversing capabilities of LRAs have been extensively investigated, their impact on immune effector cells, particularly natural killer (NK) cells, has received little attention (4–7).

The functionality of NK cells is influenced by several receptors including NKG2A and NKG2C. The ligand for these receptors is human leukocyte antigen (HLA)-E, a non-classical major histocompatibility complex (MHC) class Ib molecule that is widely expressed but at lower levels compared to classical HLA molecules. The affinity of HLA-E for NKG2A is up to six times higher than for NKG2C, meaning the inhibiting signal tends to prevail. Additionally, HLA-E expression is often increased on infected CD4^+^ T cells (8,9). Consequently, blocking the interaction between NKG2A and HLA-E could potentially unleash the cytotoxic capacity of NK cells. Indeed, monalizumab, a humanized monoclonal anti-NKG2A antibody, has demonstrated significant improvement in NK cell-mediated destruction of various tumor cell types (10). Moreover, reducing HLA-E expression on tumor cells using proteasome inhibitors has yielded positive results (11). The proteasome pathway is responsible for degrading misfolded or damaged proteins, and inhibiting this pathway leads to the accumulation of unfolded proteins in endoplasmatic reticulum (ER) which triggers ER stress. Beyond its cytotoxic effect, proteasome inhibition is thought to increase the susceptibility of multiple myeloma cells to NK-cell mediated killing by increasing the expression of MHC class I chain-related protein A/B (MICA/B) and decreasing the expression of certain other tumor cell markers (12,13). Bortezomib (Velcade®) was the first FDA-approved proteasome inhibitor, in 2003 followed by carfilzomib (Kyprolis®). These inhibitors target the 20S subunit of the proteasome, with bortezomib binding reversibly and carfilzomib binding irreversibly. While most research on proteasome inhibition has focused on hematological malignancies, its effects on HIV-1 and HIV-1 latency have been explored to a much lesser extent. Krishnan *et al.* conducted an extensive gene expression analysis of latently infected cells and discovered that genes associated with the proteasome were upregulated, suggesting that proteasome inhibition could play a role in HIV-1 latency (16). Other studies indicate that proteasome inhibition disrupts the viral replication cycle by causing an accumulation of viral proteins (14,15). Indeed, Covino *et al*. found that while bortezomib has limited capacity to reverse latency *in vitro*, it enhances the effects of romidepsin, vorinostat and panobinostat (5). Co-treatment with bortezomib and these agents led to a significant increase in NKG2D ligands MICA/B compared to single treatment. However, the impact on primary NK cells and their cytotoxic activity has not yet been explored.

In this study, we aimed to explore the impact of proteasome inhibition on latency reversal, as well as its effects on the phenotype and functionality of primary NK cells. Furthermore, we evaluated the ability of NK cells to eliminate reactivated reservoir cells by NK cells *ex vivo*.

## Results

### Inhibition of the proteasome induces latency reversal

To investigate the characteristics of HIV-1 latency, various *in vitro* latency models have been developed, including different J-Lat clones with Env/Nef-deficient pro-virus and ACH-2 cells with full-length HIV-1. These models differ in their provirus integration sites, which may affect their responsiveness to LRAs (17). We first assessed the impact of different proteasome inhibitors on latency reversal and viability in J-Lat and the ACH-2 cell lines. Bortezomib and lactacystin did not affect cell viability, while carfilzomib caused a moderate yet significant reduction in viability of J-Lat 8.4, J-Lat 10.6 and ACH-2 cells compared to DMSO-treated controls (Supplementary Figure 1a). J-Lat 6.3, J-Lat 8.4 and J-Lat 15.4 clones showed no response to the PI’s, whereas the J-Lat 10.6 and ACH-2 cells demonstrated increased sensitivity to PI-induced latency reversal (Supplementary Figure 1b). Based on these results, we chose to proceed with the J-Lat 8.4 (insensitive) and J-Lat 10.6 (sensitive) clone for further experiments. We will focus on bortezomib, the most extensively studied PI.

Next, we investigated whether bortezomib could augment latency reversal when used in combination with other LRAs. Romidepsin, vorinostat and TNF-α all effectively induced latency reversal in the J-Lat 8.4 cell line, while the SMAC mimetic birinapant did not (Figure 1a-b). In the J-Lat 10.6 cell line, significant latency reversal was observed with all individual treatments (Figure 1c-d). The combination of bortezomib and vorinostat resulted in significantly greater latency reversal compared to vorinostat alone. However, combining bortezomib with either romidepsin, TNF-α or birinapant did not produce greater latency reversal than was achieved with single LRA treatment (Figure 1c-d). Although none of the tested LRAs affected the viability of J-Lat 8.4 and J-Lat 10.6 cells (Supplementary Figure 2), the viability was significantly decreased when bortezomib was combined with romidepsin, TNF-α, vorinostat or birinapant,.

**Figure 1.**
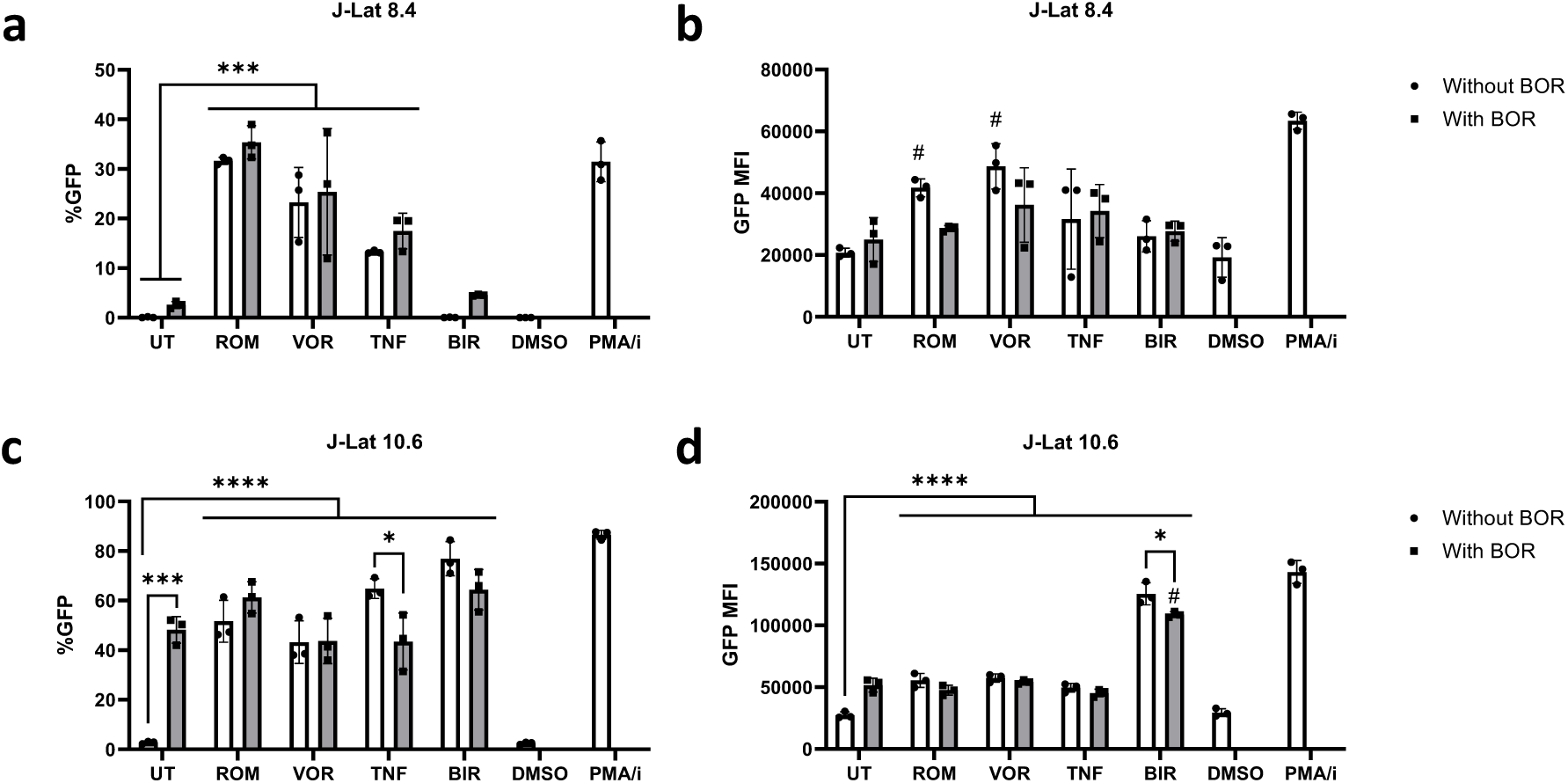
Effect of bortezomib on latency reversal in J-Lat cells. J-Lat 8.4 cells and J-Lat 10.6 cells were treated for 24h with either 10 nM romidepsin (ROM), 5 µM vorinostat (VOR), 10 ng/mL TNF-α, 50 nM birinapant (BIR) alone or in combination with 10 nM bortezomib (BOR). DMSO and PMA/I were used as negative and positive control, respectively. **a)** Percentage of latency reversal in J-Lat 8.4 cells measured by GFP and **b)** GFP MFI (n =3; mean ± SD). **c)** Percentage of latency reversal in J-Lat 10.6 cells measured by GFP and **d)** GFP MFI (n =3; mean ± SD). Two-way ANOVA with Tukey’s multiple comparisons test. Significance levels were graphically annotated as follows: * p<0.05; ** p<0.01 and ***p<0.001. Statistically significant differences compared to untreated condition are indicated with #.

### Bortezomib reduces HLA-E expression in reactivated J-Lat cells

Since the impact of proteasome inhibition on HLA-E expression has not been explored in the context of HIV-1 latency, we aimed to address this question. All tested LRAs, with the exception of TNF-α, reduced HLA-E expression compared to untreated J-Lat 8.4 and J-Lat 10.6 cells. Additionally, treatment with bortezomib further decreased HLA-E expression compared to LRA treatment alone. This reduction in HLA-E expression was notably more pronounced in J-Lat 8.4 cells compared to J-Lat 10.6 cells (Figure 2 and Supplementary Figure 3). Following single LRA treatment, the percentage of GFP^+^HLA-E^+^ J-Lat cells was higher compared to the percentage of GFP^+^HLA-E^-^ J-Lat cells. However, combining romidepsin or vorinostat with bortezomib significantly increased the proportion of GFP^+^HLA-E^-^ J-Lat cells. In contrast, bortezomib did not increase the proportion of GFP^+^HLA-E^-^ J-Lat cells when combined with TNF-α or birinapant (Figure 2c and f). Furthermore, bortezomib treatment did not affect the expression of HLA class I molecules on J-Lat 8.4 and J-Lat 10.6 cells (Supplementary Figure 4).

**Figure 2.**
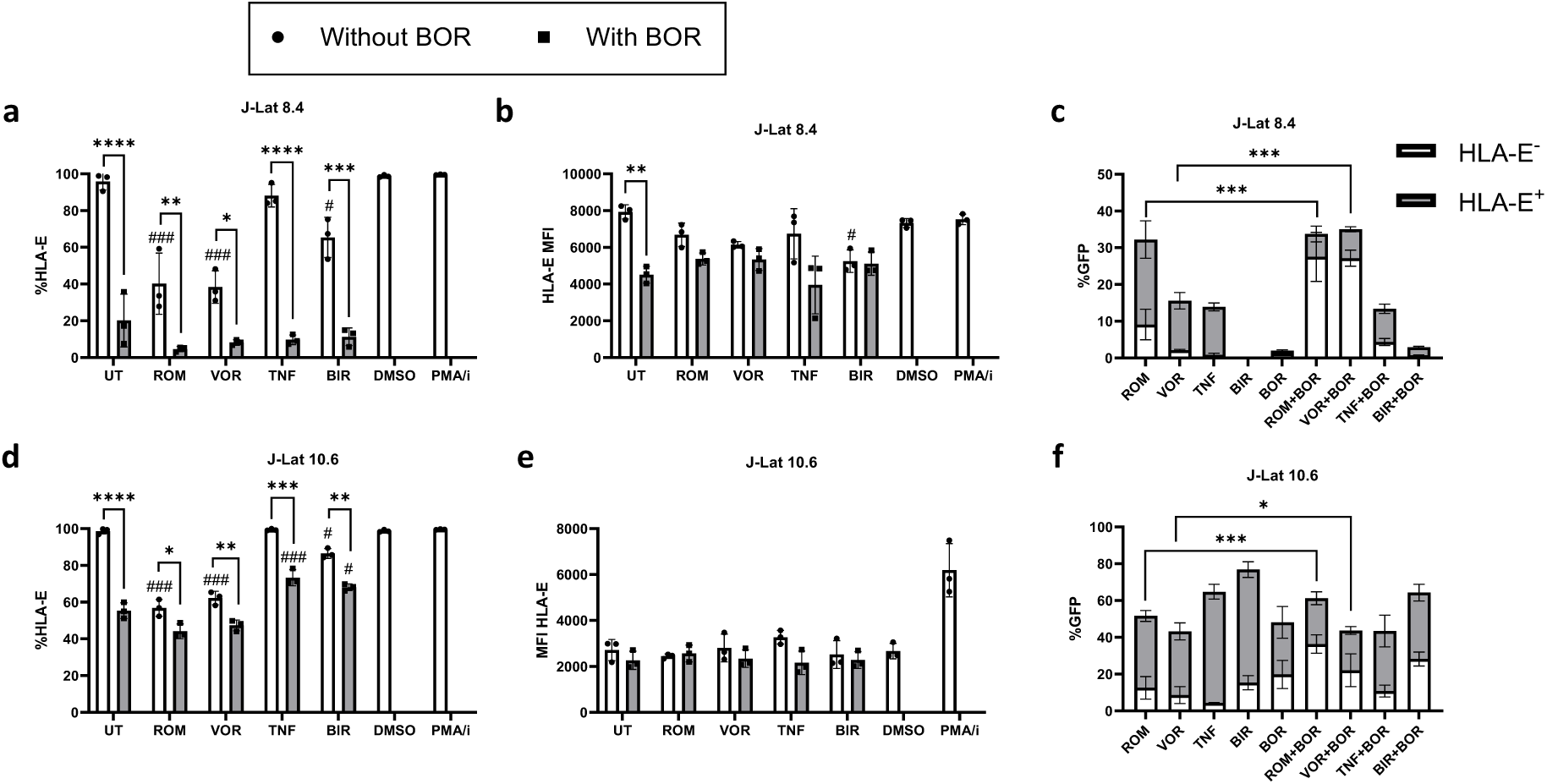
Effect of bortezomib on HLA-E expression in J-Lat cells. J-Lat 8.4 cells or J-Lat 10.6 cells were treated overnight with indicated LRAs alone or in combination with 10 nM bortezomib (BOR). a) Percentage of HLA-E expressing J-Lat 8.4 cells and b) respective MFI of HLA-E. c) Percentage of reactivated J-Lat 8.4 cells expressing HLA-E or not. d) Percentage of HLA-E expressing J-Lat 10.6 cells and e) respective MFI of HLA-E. f) Percentage of reactivated J-Lat 10.6 cells expressing HLA-E (gray) or not (white). Two-way ANOVA with Tukey’s multiple comparisons test. Significance levels were graphically annotated as follows: * p<0.05; ** p<0.01; ***p<0.001 and **** p<0.0001. Statistically significant differences compared to untreated condition are indicated with #.

To explore the mechanism by which bortezomib reduces HLA-E expression, we treated K562 cells stably transduced with HLA-E with bortezomib and observed a decrease in HLA-E levels, indicating a post-translational mechanism rather than a transcriptional effect (Supplementary Figure 5). Pre-treating K562*HLA-E or J-Lat 8.4 cells with cycloheximide (CHX), an antibiotic that inhibits the elongation step of the translation process, did not significantly alter HLA-E expression between bortezomib treated and bortezomib/CHX treated cells. This suggests that bortezomib reduces HLA-E expression through a mechanism independent of translation, potentially involving ER-stress induced misfolding of HLA-E proteins.

### Effect of bortezomib on MICA/B expression on J-Lat cells

Next, we evaluated the impact of LRA and/or bortezomib treatment on the expression of NKG2D ligands, MIC-A and MIC-B. In both J-Lat cell lines, treatment with bortezomib, TNF-α and birinapant did not significantly alter MIC-A and MIC-B expression. As anticipated, romidepsin and vorinostat significantly increased the proportion of MIC-A and MIC-B expressing J-Lat cells (Supplementary Figure 6). Single treatment with vorinostat significantly decreased the percentage of NKG2D-expressing NK cells, whereas bortezomib, whether used alone or in combination, did not affect NKG2D expression on NK cells (Supplementary Figure 7).

### Bortezomib increases the proportion of cytokine producing NK cells

The effects of LRAs on NK cells and other immune cells have been minimally studied (6,18) and to our knowledge, the impact of bortezomib on NK cells remains unexplored. Therefore, we examined how LRAs, with or without bortezomib, affect the phenotype of primary NK cells (Figure 3a). Based on the expression of CD56 and CD16, four different NK cell subsets were distinguished, as previously described by our group (19). CD56^br^ NK cells are immature and differentiate into cytokine-producing CD56^dim^CD16^-^ NK cells and cytotoxic CD56^dim^CD16^+^ NK cells. Treatment with LRAs did not significantly impact the viability of primary NK cells while bortezomib significantly decreased their viability. However, combining bortezomib with LRAs did not further affect NK cell viability compared to single treatments (Figure 3b).

**Figure 3.**
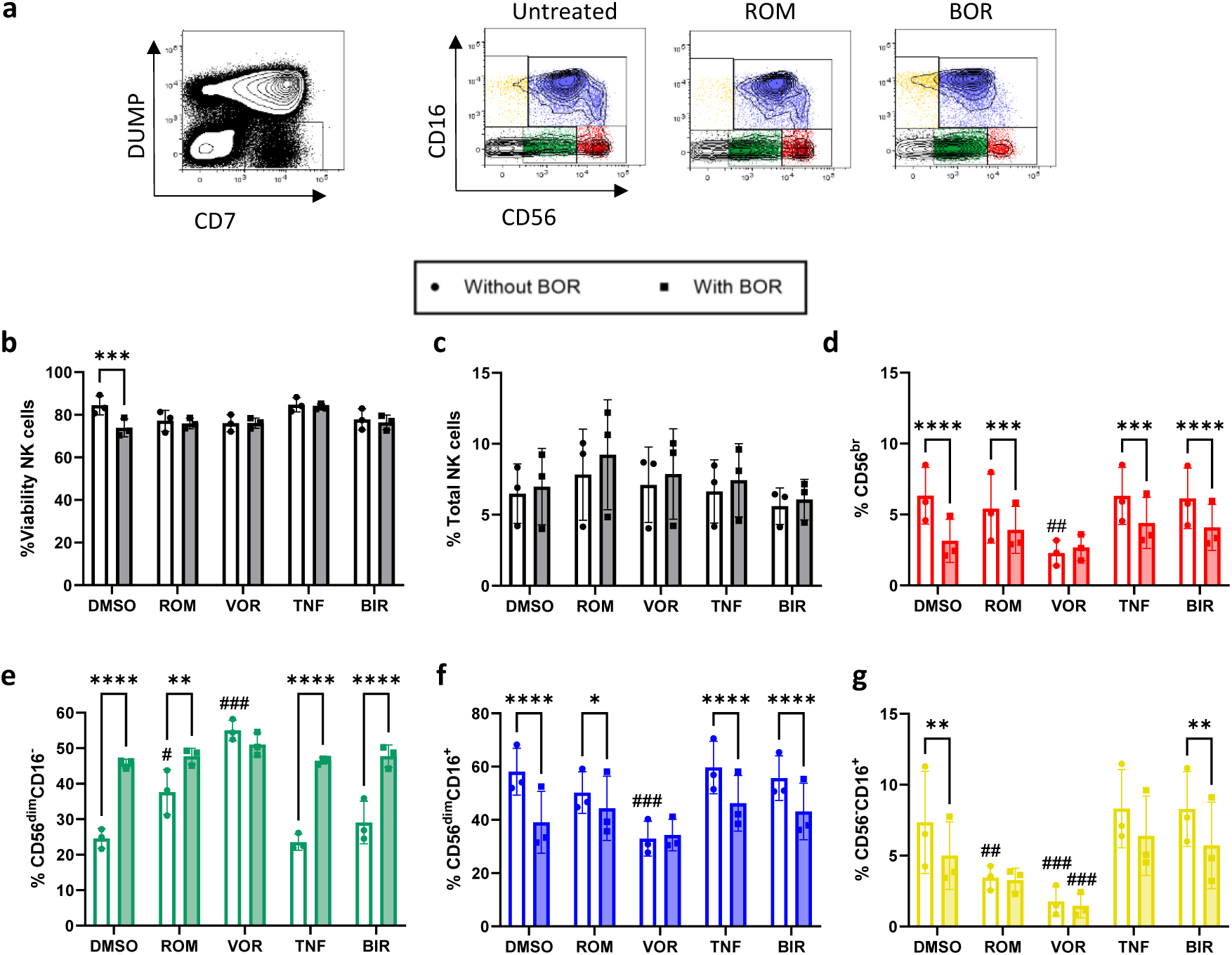
Effect of bortezomib on viability and frequency of primary NK cells. PBMCs from healthy donors were treated for 24 hours with either LRA alone (empty bars) or in combination with bortezomib (BOR, filled bars). a) Gating strategy for NK cells and NK cell subsets. NK cells were identified based on the absence of CD3, CD14 and CD19 (DUMP) and the presence of CD7. NK cell were subsequently divided into subsets based on CD56 and CD16 expression. b) Viability of NK cells was assessed using FVS450. c) Total NK-cell frequency measured by flow cytometry as DUMP^-^CD7^+^ cells. d-g) Frequency of NK cell subsets (according to CD56 and CD16 expression). Two-way ANOVA with Tukey’s multiple comparisons test. Significance levels were graphically annotated as follows: * p<0.05; ** p<0.01; ***p<0.001 and **** p<0.0001. Statistically significant differences compared to untreated condition are indicated with #.

The overall frequency of primary NK cells did not significantly change with treatment by single LRAs or in combination with bortezomib (Figure 3c). However, there was a significant decrease in the proportion of cytokine-producing CD56^bright^ and cytotoxic CD56^dim^CD16^+^ NK cells following single bortezomib treatment and in combination with romidepsin, TNF-α or birinapant (Figure 3d and f). In addition, except for vorinostat, the frequency of cytokine-producing CD56^dim^CD16^-^ NK cells significantly increased when bortezomib was added to LRAs. Single treatment with vorinostat already caused a significant increase in CD56^dim^CD16^-^ NK cells while decreasing the other three NK-cell subsets (Figure 3e). Compared to untreated NK cells, the dysfunctional CD56^-^CD16^+^ subset significantly decreased with treatments of bortezomib, romidepsin and vorinostat (Figure 3g).

### Effect of bortezomib on the expression of NK cell receptors

Given that bortezomib reduces HLA-E expression on J-Lat 8.4 and J-Lat 10.6 cells, we next investigated its effect on the expression of the NK cell receptors that recognize HLA-E. Initially, we assessed bortezomib’s impact on NKG2A expression on the NK-92 cell line and observed a dose-dependent reduction in NKG2A expression (Supplementary Figure 8). We then explored how LRAs and/or bortezomib affect NKG2A/C expression on primary NK cells. Romidepsin increased NKG2A expression, whereas vorinostat, TNF-α and birinapant significantly decreased NKG2A expression compared to untreated NK cells, both in terms of proportion (% positive) and on a per cell basis (MFI) (Figure 4b). Addition of bortezomib further diminished NKG2A expression. In contrast, the expression of NKG2C on primary NK cells remained unchanged with LRAs and/or bortezomib (Figure 4a and b). Bortezomib, TNF-α and birinapant, when used individually, reduced the CD94 expression on NK cells, as indicated by a significantly lower MFI. However, combining bortezomib with LRAs did not further decrease CD94 expression (Figure 4a and b).

**Figure 4.**
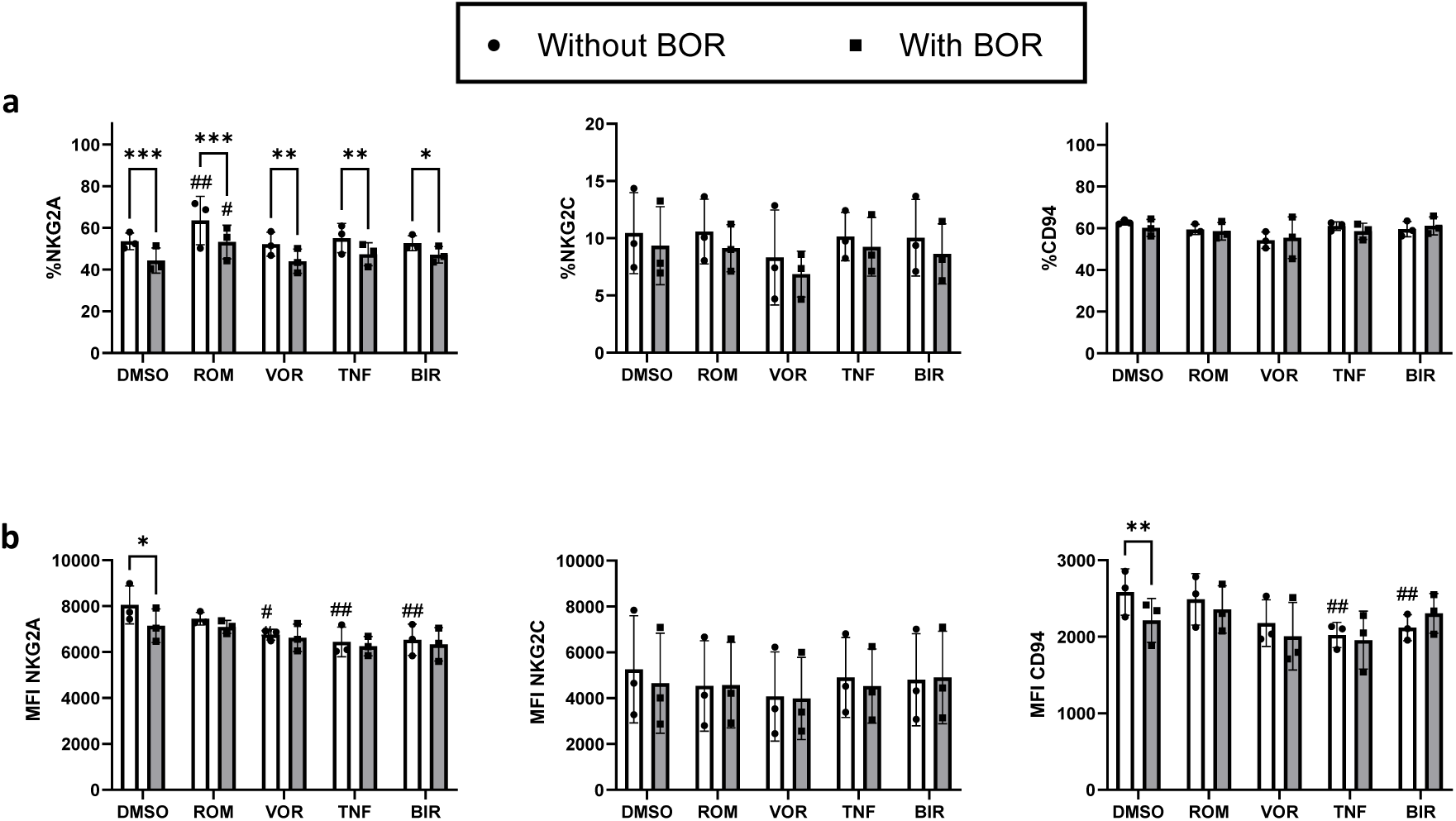
Effect of bortezomib on the expression of HLA-E receptors on NK cells. PBMCs were treated for 24h with either LRA alone or in combination with bortezomib (BOR). Percentage of primary NK cells (DUMP^-^CD7^+^ cells) expressing a) NKG2A, b) NKG2C or c) CD94. Two-way ANOVA with Tukey’s multiple comparisons test. Significance levels were graphically annotated as follows: * p<0.05; ** p<0.01; ***p<0.001 and **** p<0.0001. Statistically significant differences compared to untreated conditions are indicated with #.

Next, we assessed the expression of the NKG2A/C receptors across different NK-cell subsets (Supplementary Figure 9). We found that the percentage of NKG2A and CD94-expressing NK cells was significantly reduced in the cytotoxic CD56^dim^CD16^+^ NK cell subset across all treatment conditions. Bortezomib alone increased the proportion of NKG2C-expressing NK cells only in the cytokine-producing CD56^br^ subset. Similarly, single treatments with romidepsin and vorinostat also elevated the proportion of NKG2C expressing CD56^br^ NK cells. However, when bortezomib was combined with romidepsin, NKG2C expression was decreased in both CD56^dim^CD16^+^ (cytotoxic) and CD56^dim^CD16^-^ (cytokine-producing) subsets, compared to single LRA treatments.

### Effect of bortezomib on direct NK cell cytotoxicity

Additionally, we investigated the effect of combining bortezomib with LRA treatment on NK-cell cytotoxicity against K562 cells devoid of MHC. Bortezomib alone did not influence NK cell-mediated lysis of K562 cells or NK cell degranulation compared to DMSO-treated NK cells. Among the tested LRAs, only TNF-α significantly increased NK cell degranulation towards K562 cells in both PLWH and healthy donors. However, this increase in degranulation did not translate into enhanced lysis of K562 cells. The combination of bortezomib with TNF-α led to a significant reduction in NK cell-mediated lysis of K562 cells in PLWH, but not in healthy donors (Figure 5a). Nevertheless, bortezomib significantly enhanced the degranulation of primary NK cells only when combined with TNF-α during these co-cultures (Figure 5c).

**Figure 5.**
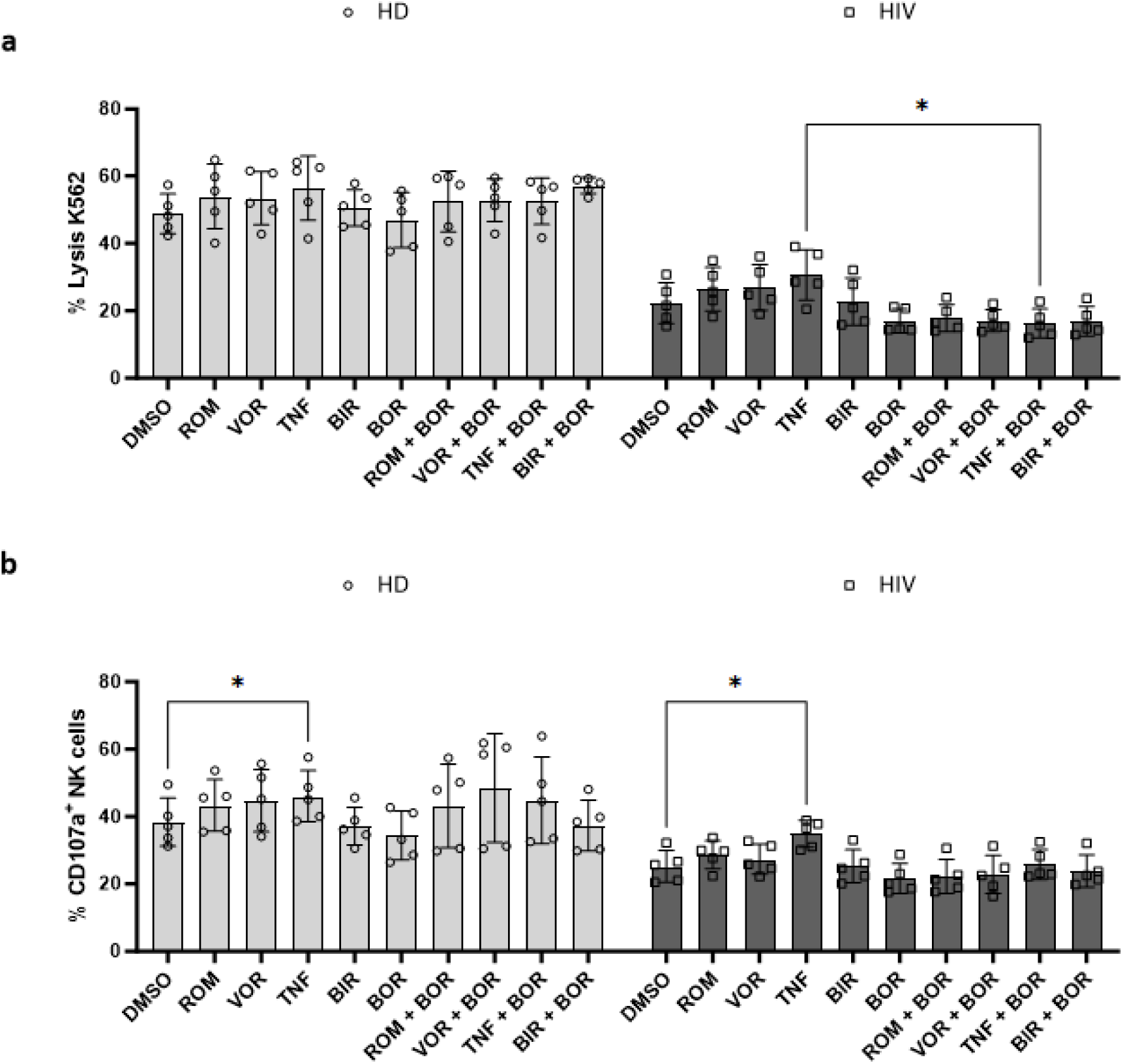
Effect of bortezomib on direct NK cell mediated cytotoxicity. PBMCs from healthy donors or PLWH were treated overnight with either LRA alone or in combination with bortezomib. Subsequent co-cultures with MHC-devoid K562 cells were performed (6h). a) Specific lysis of K562 target cells was measured using FVS780. b) Degranulation of primary NK cells was measured using CD107a. Two-way ANOVA with Tukey’s multiple comparisons test. Significance levels were graphically annotated as follows: * p<0.05; ** p<0.01; ***p<0.001 and **** p<0.0001. Statistically significant differences compared to untreated (blue) or single BOR treatment condition are annotated with #.

### NK cell-mediated killing of reactivated J-Lat cells

Since bortezomib impacts both the expression of NKG2A/C receptors on NK cells and HLA-E on target cells, we examined whether this would enhance NK-cell functionality. For this, we co-cultured reactivated J-Lat 8.4 and J-Lat 10.6 cells with primary NK cells from healthy donors or PLWH. Unlike single cultured J-Lat 8.4 cells, bortezomib and all tested LRAs except TNF-α, did not enhance NK cell mediated killing of reactivated J-Lat 8.4 cells (Supplementary Figure 10 and 11a). In fact, combining romidepsin with bortezomib even reduced the NK-cell mediated elimination of these cells. Conversely, TNF-α, birinapant, bortezomib and the combination of romidepsin and bortezomib significantly increased cell death in J-Lat 10.6 cells (Figure 6b). Interestingly, the percentage of GFP expressing J-Lat 8.4 and J-Lat 10.6 cells significantly decreased with LRA treatments and in combination with bortezomib (Figure 6b and Supplementary Figure 10b). This effect was further enhanced by the anti-NKG2A antibody, monalizumab, in romidepsin-treated J-Lat 10.6 cells indicating an NKG2A-mediated mechanism of J-Lat elimination by NK cells (Supplementary Figure 12). However, this effect was less pronounced in bortezomib and romidepsin/bortezomib-treated J-Lat 10.6 cells likely due to the bortezomib-mediated decrease in HLA-E expression. In contrast, treatment with a blocking anti-NKG2C antibody did not affect GFP^+^ J-Lat 10.6 cell levels or cell death.

**Figure 6.**
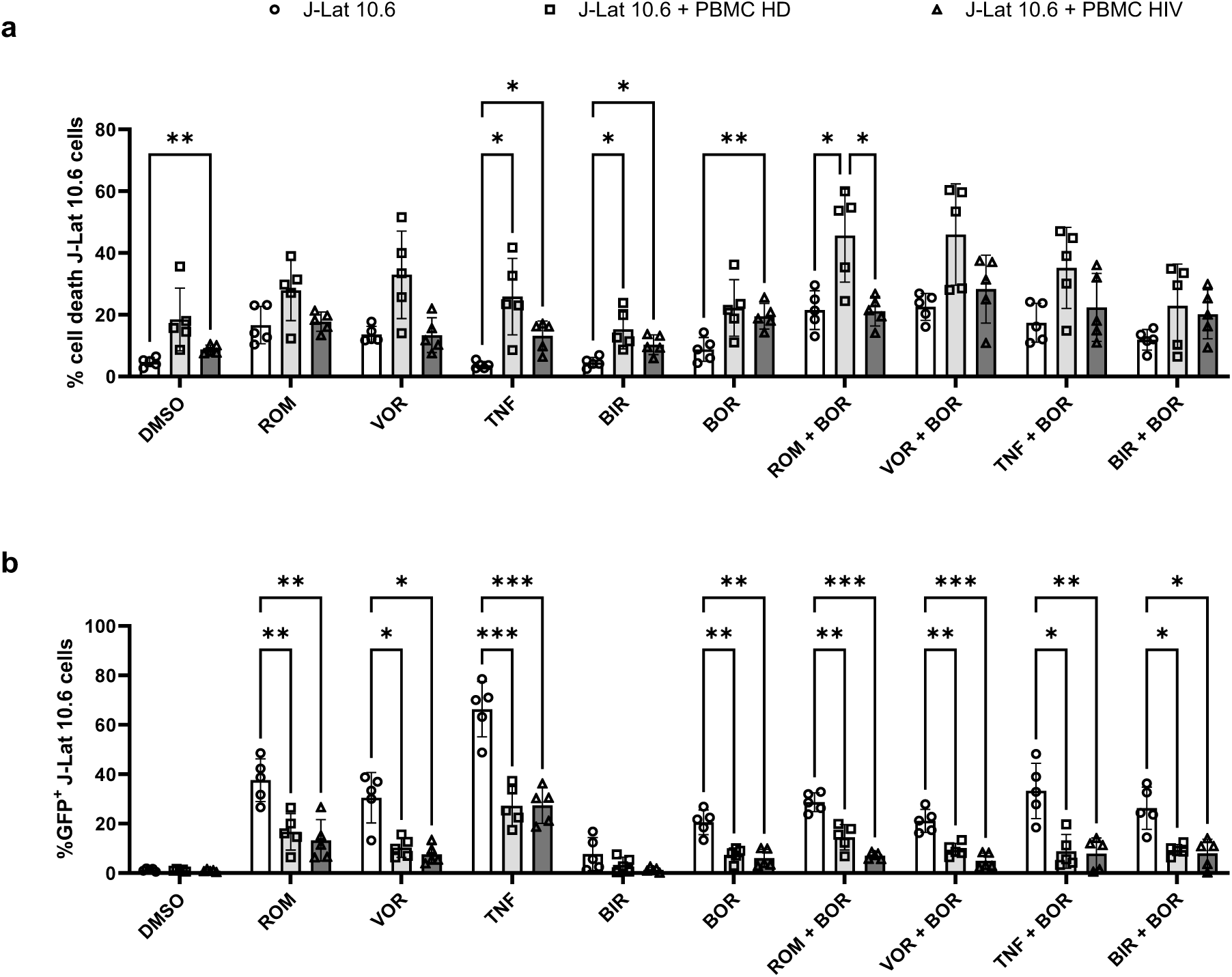
Effect of bortezomib on NK cell mediated killing of reactivated J-Lat 10.6 cells. a) Viability of J-Lat 10.6 cells after reactivation and subsequent co-culture with PBMCs, measured by FVS780. B) GFP expression in J-Lat 8.4 cells after reactivation and subsequent co-culture with PBMCs. Two-way ANOVA with Tukey’s multiple comparisons test. Significance levels were graphically annotated as follows: * p<0.05; ** p<0.01; ***p<0.001 and **** p<0.0001. Statistically significant differences compared to single LRA condition are annotated with #.

## Discussion

In the present study, we explored the effects of adding bortezomib to LRA treatment on HIV-1 reactivation, HLA-E expression and subsequent elimination of reactivated cells by NK cells *in vitro*.

We initially demonstrated that bortezomib alone can induce latency reversal in J-Lat 10.6 cells while other J-Lat clones did not respond similarly. This variance might be due to differences in proviral integration sites between the latent cell lines. Additionally, bortezomib did not enhance the latency-reversing capacity of HDACi, TNF-α or birinapant in J-Lat 8.4 and J-Lat 10.6 cell lines. Conversely, Covino *et al.* observed a synergistic effect on latency reversal in J1.1 cells when bortezomib was combined with vorinostat or romidepsin, while bortezomib alone did not influence latency reversal (5). Given that similar concentrations of LRAs and bortezomib were used in the latter study compared to ours, the differing results are likely due to the distinct genetic backgrounds of the cell lines used. Therefore, these findings should be validated on primary cells as well as other cellular and anatomical reservoirs. Cary *et al.* reported that bortezomib could significantly induce latency reversal at very low concentrations (0.5 nM) in latently infected primary CD4^+^ T cells via activation of the NF-κB pathway (20). Notably, adding bortezomib to vorinostat did not produce a synergistic latency reversal effect in their study. Importantly, bortezomib does not induce T-cell activation making it an ideal LRA candidate (21).

Furthermore, our study demonstrated that bortezomib reduces HLA-E expression on J-Lat 8.4 and J-Lat 10.6 cells without affecting HLA class I expression, consistent with findings in MM. The proposed mechanism for this effect involves proteasome inhibition by bortezomib, leading to ER-stress, which subsequently causes improper peptide loading on HLA-E in the ER and reduced translocation to the cell membrane. The differential effect on HLA-E and HLA class I expression may stem from HLA-E’s greater reliance on accurate peptide loading for its expression and stability compared to other HLA class I molecules (22).

Additionally, we observed a notable decrease in HLA-E expression following individual treatments with romidepsin and vorinostat. This is in contrast with the findings of Garrido *et al.* who reported only a slight and statistically insignificant decrease in HLA-E expression (23). Notably, our study revealed that the reduction in HLA-E expression primarily occurs in J-Lat cells emerging from latency, which may have important implications for NK cell-mediated clearance of reactivated cells.

While cytotoxic T lymphocyte (CTL) responses are generally considered more effective than NK-cell responses during the natural course of HIV infection, our study focussed on NK-cell mediated clearance of reactivated cells. NK cells are thought to play a more critical role in eliminating reactivated cells compared to CTLs (24). This is particularly important because most latent proviruses contain escape mutations that allow them to evade CTL responses, but these mutations do not affect NK-cell activity (25). Additionally, in PLWH a reduction in viral DNA following treatment with panobinostat was associated with enhanced NK-cell responses rather than HIV-1 specific CTL responses (26).

It is essential for LRAs to avoid negatively affecting effector cells, including NK cells, as the efficient elimination of reactivated infected cells by these effector cells is critical. In our study, the viability of primary NK cells after 24h of bortezomib treatment was only slightly affected, aligning with previous findings. However, it is important to note that most studies evaluate bortezomib toxicity over a longer period, up to 72h, when adverse effects become more pronounced (5).

Additionally, we observed no changes in the overall frequency of NK cells with any treatment. Nevertheless, there was a significant decline in the number of cytotoxic NK cells accompanied by increase in the number of cytokine-producing NK cells. The reduction in CD16 expression, previously reported for treatments like prostratin or bryostatin, has been attributed to matrix metalloprotease-mediated shedding of CD16 (6). Such downregulation of CD16 may impair ADCC-mediated elimination of target cells. Indeed, it has been shown that pre-treatment of primary NK cells with bryostatin but not prostratin inhibits the killing of rituximab-coated Raji cells (6). However, this specific aspect was not evaluated in our study.

While the expression of NKG2D was only minimally affected by bortezomib treatment, significant differences in the expression of NK cell receptors for HLA-E were observed. Combined LRA and bortezomib treatment resulted in a significant decrease in NKG2A and CD94 expression in the cytotoxic NK cell subset but not in other subsets. Conversely, the expression of NKG2C showed distinct responses depending on the NK cell subtype. Whereas the CD56^br^ subset showed an increased expression of NKG2C, the opposite was observed for CD56^dim^ subset.

The bortezomib-induced reduction in HLA-E expression on reactivated J-Lat cells together with the decrease of NKG2A on NK cells could have important implications for NK-cell cytotoxicity and overall functionality. Our results suggest that adding bortezomib to LRA treatment enhances both direct cytotoxicity against K562 cells and against reactivated J-Lat cells. However, inconsistent findings were noted regarding the reduction in viability of J-Lat cells and the decrease in GFP expression in co-cultures between PBMCs and reactivated J-Lat cells. Enhancing the LRA-bortezomib combination with IL-15 might further amplify NK-cell responses against reactivated reservoir cells.

Therapeutic disruption of the HLA-E/NKG2A axis using monalizumab has shown promising results, but it may lead to general hyporesponsiveness in NK cells and CD8^+^ T cells. Therefore, we suggest a more targeted strategy that involves specific downregulation of HLA-E on target cells, such as latently infected cells or tumor cells, as a more favourable approach. Previous clinical trials involved treating multiple myeloma patients with bortezomib followed by NK-cell transfusions after a washout period. However, our findings indicate that bortezomib does not negatively affect NK-cell functionality, suggesting that a washout period between bortezomib infusions and NK-cell injections may not be necessary.

Although bortezomib is commonly used to treat certain malignancies, it is associated with significant toxicities, including neurological issues. However, newer generations of proteasome inhibitors, such as carfilzomib and ixazomib, offer similar effectiveness with fewer side effects. In this study, we compared the latency reversing abilities and HLA-E downregulation effects of bortezomib, carfilzomib and lactacystin and found them to be comparable. Reid *et al*. also explored the use of bortezomib in patients with AIDS-associated Kaposi sarcoma and reported a significant reduction in pVL, potentially due bortezomib-induced increases in APOBEC3G levels, which impair the viral replication cycle (27).

Several important limitations of this study should be acknowledged. First, our research was confined to the blood compartment excluding other tissues such as GALT, which are known contributors to the viral reservoir. Second, our *in vitro* analysis employed single doses of LRAs and bortezomib, which differs from *in vivo* clinical trials where multiple doses are typically administered over time. Additionally, potential interactions between bortezomib/LRA combinations and ART were not addressed and warrant further investigation.

In conclusion, our study reveals that bortezomib not only reduces HLA-E expression on reactivated reservoir cells, but also alters the expression of the HLA-E receptor, NKG2A, on NK cells, leading to enhanced NK-cell mediated elimination of reactivated cells *in vitro* (Figure 7, graphical abstract).

**Figure 7.**
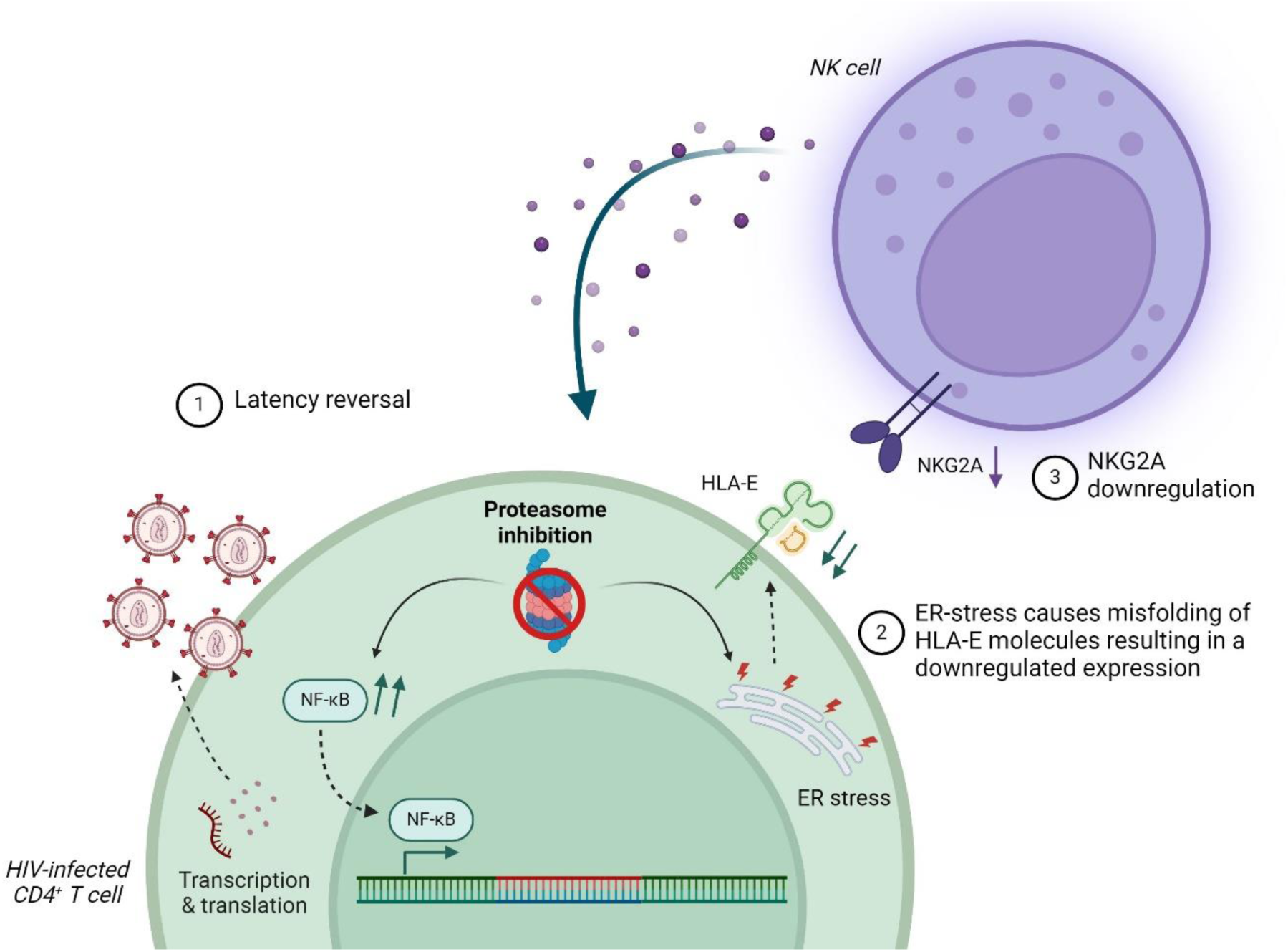
Graphical abstract. Inhibition of the proteasome leads to latency reversal through several mechanisms: 1) It elevates the intranuclear levels of the transcription factor NF-κB, 2) It reduces HLA-E expression by causing ER-stress mediated misfolding of HLA-E molecules and 3) It downregulates NKG2A on primary NK cells via an unspecified mechanism. Figure created with BioRender.com.

## Methods

### Primary cells and cell lines

Blood samples were obtained from healthy volunteers and PLWH after receiving informed consent (Ethical Committee of UZ Brussels, ref. 2019/247). Peripheral blood mononuclear cells (PBMCs) were isolated using Lymphoprep density gradient centrifugation. Isolated PBMCs were frozen in Cryostor (Stemcell Technologies, Vancouver, Canada) and stored in liquid nitrogen until further use. Upon thawing, PBMCs were cultured overnight in Iscove’s modified Dulbecco medium (IMDM, Lonza, Basel Switzerland) containing 10% fetal bovine serum (FBS, Tico Europe, Amstelveen, The Netherlands), 2 mM L-glutamine (Sigma-Aldrich, Ghent, Belgium) and 100 international units (IU)/mL streptomycin, 100 µg/mL penicillin (Sigma-Aldrich) and 20 IU/mL recombinant human IL-2 (rhIL-2, Proleukin, Clinigen, Trent, United Kingdom), referred to as complete IMDM.

J-Lat 8.4 cells (generously provided by Prof. Rob Gruters, Virosciences, Erasmus Medical Center Rotterdam, the Netherlands), J-Lat 6.3, J-Lat 10.6, J-Lat 15.4, A3.01 and ACH-2 cells (kindly provided by Prof. Carine Van Lint, ULB, Belgium), Jurkat and K562 cells (kindly provided by Prof. Karine Breckpot, VUB) were cultured in Roswell Park Memorial Institute (RPMI)-1640 medium (Lonza) supplemented with 10% FBS, 2 mM L-glutamine, 100 IU/mL Streptomycin and 100 µg/mL penicillin (complete RPMI). K562 cells expressing HLA-E (K562*HLA-E, kindly provided by Prof. Thorbald van Hall, Medical Oncology, Leiden University) were cultured in complete RPMI containing 2 µg/mL blasticidin (Sigma-Aldrich). NK-92 cells (courtesy of Prof. Karine Breckpot, Laboratory of Molecular and Cellular Therapies, VUB) were maintained in alpha-Minimum Essential Medium (α-MEM, Sigma-Aldrich) supplemented with 12.5% FBS, 12.5% horse serum (Invitrogen, Massachusetts, USA), 2 mM L-glutamine, 200 U/mL recombinant human IL-2 (Proleukin). All cells were incubated at 37°C with 5% CO_2_ in a humidified incubator.

### Flow cytometry

Cell viability was assessed using Fixable viability Dye eFluor780 (Thermo Fisher, 1/3500 dilution in dPBS). The staining process was conducted at room temperature for 20 minutes in the dark followed by washing the cells with dPBS. For labelling cell surface molecules, cells were incubated with specific mAbs (see Supplementary Table 1) for 30 minutes at 4°C in the dark, using FACS buffer (dPBS containing 1% bovine serum albumin (BSA) and 0.1% sodium azide) as the diluent. Primary NK cells were identified as CD7^+^ cells after exclusion of T cells (CD3), B cells (CD19) and monocytes (CD14) (DUMP^-^). Classification of NK cell subsets was based on CD16 and CD56 expression. Data acquisition was performed using the BD LSR Fortessa.

### Latency reversal and HLA-E assessment

J-Lat 8.4 or J-Lat 10.6 cells (150,000 cells) were cultured in a 24-well plate with complete RPMI supplemented with the indicated concentrations of LRAs. The LRAs used included: 10 nM romidepsin (Selleckchem, Houston, USA), 10 µM vorinostat (Selleckchem), 10 ng/mL TNF-α (Immunotools, Friesoythe, Germany), 50 nM birinapant (Selleckchem), 10 nM bortezomib (Selleckchem), 40 nM carfilzomib (Selleckchem) and 20 µM lactacystin (Cayman Chemical Company, Michigan, USA). The highest concentration of DMSO (0.2%) served as a negative control, while 20 ng/mL PMA with 500 ng/mL ionomycin was used as a positive control. Latency reversal was assessed by measuring GFP expression after 24h of incubation. HLA-E expression was assessed on J-Lat cells and K562*HLA-E cells after labelling with specific fluorescent antibodies.

### NK cell cytotoxicity assay

PMBCs were thawed and allowed to rest overnight in complete IMDM at a density of 2×10^6^ cells/mL in a 37°C, 5% CO_2_ humidified incubator. The following day, PBMCs were treated overnight with the indicated LRAs, with or without bortezomib. Co-cultures with wild-type K562 cells (targets) were set up in a 96 U-bottom well plate at a 10:1 effector to target (E:T) ratio for 6 hours. After one hour, 5 µg/mL Brefeldin A, 2 µM Monensin (BioLegend, San Diego, CA, USA) and CD107a (1 µL/well) were added to each well. Cell death was assessed using FVS eFluor780.

### NK cell-mediated killing of reactivated J-Lat cells

J-Lat 8.4 or J-Lat 10.6 cells were cultured overnight (16h-18h) in a 96-well U-bottom plate with complete RPMI containing the indicated concentrations of LRAs. Meanwhile, PMBCs were thawed and rested overnight in complete IMDM at a density of 2×10^6^ cells/mL in a 37°C, 5% CO_2_ humidified incubator. The following day, PBMCs were stained with 0.5 µM CellTrace^TM^Violet (Thermo Fisher) to discriminate between J-Lat cells and PBMCs. Reactivated J-Lat cells (targets) were then seeded in a U-bottom 96-well plate along with primary PBMCs in the presence or absence of 10 µg/mL anti-NKG2A (monalizumab, MedChem Express) or anti-NKG2C (R&D Systems) for 6 hours, with or without primary NK cells (5:1 ratio). Cell death and latency reversal in J-Lat cells was assessed using FVS780 and GFP, respectively.

### Statistical analysis

Flow cytometric analysis was performed using Flowlogic v7 (Inivai, Australia). Graphs were created, and statistical analysis was performed using GraphPad Prism 10.0.0 (San Diego, CA, USA). Data are presented as mean ± standard deviation (SD). Unless indicated otherwise in the figure legend, statistical comparisons were made using a two-way ANOVA with Sidak’s correction for multiple comparisons. P values of <0.05 were considered statistically significant. Significance levels were graphically annotated as follows: * p<0.05; ** p<0.01; ***p<0.001 and **** p<0.0001.

## Data availability

All data generated during this study are included in the article and its Supplementary Information. The data that support the finding of the current study are available from the corresponding author upon request.

## Acknowledgments

We would like to thank all study participants for their collaboration. J.L.A. and S.D.A. received funding from Scientific Fund Willy Gepts of the UZ Brussel and the Strategic Research Program (SRP089).

## Author contributions

SDA and JLA provided funding and supervised the study and were intellectually involved in the conceptualization and design of the project. TL wrote the manuscript and all co-authors revised and edited the manuscript. TL, SDR and SN contributed to the methodology. TL performed data analysis.

